# Vascular tree structure-based perfusion phantom fabrication using modified Hele-Shaw Cell technique

**DOI:** 10.64898/2026.04.29.721575

**Authors:** Susweta Das, Makrand Rakshe, Soumyajit Sarkar, Ria Paul, Shruti D. Marathe, Nixon M. Abraham, Prasanna S. Gandhi, Hari M Varma

## Abstract

Tissue phantoms that mimic microvasculature and perfusion are essential for modelling vascular function, guiding interventions, and calibrating imaging systems, which require faithful replication of vascular geometry and flow. Conventional fabrication strategies, including wire-based molding, lithographic micromachining, and additive manufacturing, offer useful capabilities but remain constrained by predefined designs, rectangular channel cross-sections, limited scalability, and high production costs. Reliance on predefined digital vascular models restricts design flexibility and limits the ability to capture the natural variability and complexity of real vascular systems. Here, we present a lithography-free, fractal-generating approach based on a modified Lifted Hele–Shaw Cell (LHSC) technique, in which vascular networks emerge spontaneously via interfacial fluid instabilities. Unlike pre-designed methods, these structures are governed by fluid properties and flow conditions, enabling adaptive, physiologically relevant geometries with smooth Gaussian cross-sections and natural diameter tapering. We demonstrate four phantom designs: a planar vascular tree, an anatomically guided cerebral network, a retinal vascular model, and a conformable curved substrate phantom. Validation using Laser Speckle Contrast Imaging confirms structural fidelity and physiologically relevant flow consistent with Murray’s law. This platform uniquely integrates realistic vascular architecture with emergent, fractal driven formation, highlighting its potential as a reproducible and biologically relevant alternative to conventional vascular phantom fabrication. Furthermore, the availability of such realistic in vitro vascular models can reduce reliance on animal experiments and contribute towards more ethical and sustainable preclinical research.

## 1 Introduction

Microvascular structures of vascular networks are essential across multiple domains in biomedical research such as disease modeling, tissue engineering, and perfusion imaging, where realistic branching geometry and perfusion dynamics are needed to replicate physiologically relevant flow conditions [1, 2]. One major application lies in perfusion imaging, which enables visualization and quantification of blood flow, allowing clinicians and researchers to study vascular disorders, guide treatments, and monitor recovery [3]. Multiple modalities are used for this purpose, including ultrasound [4], Contrast-Enhanced Magnetic Resonance Imaging (CE-MRI) [5], Computed Tomography (CT) [6], and optical methods such as Diffuse Correlation Spectroscopy (DCS) and Laser Speckle Contrast Imaging (LSCI) [7–10]. Across all these techniques, flow phantoms, which are synthetic models that replicate blood flow, play an essential role in validation and standardization.

Several strategies have been developed to fabricate vascular phantoms. In order to mimic microvasculature with circular cross-section of vascular lumen, silicon tubing of sub-millimeter diameters have been used in MRI phantoms [11]. In ultrasound imaging string-based approaches have been used to build phantoms where soldered wires are shaped into branching vessels and embedded in Tissue-Mimicking Materials (TMMs) such as gelatin, silicone, and plastisol, which are later removed to create channels [12–14]. Advancements in manufacturing techniques have further expanded phantom design possibilities. Three-dimensional printing has enabled accurate patient-specific vascular models for MRI [15] and cerebral perfusion phantoms for CT [16].

More precise microfabrication approaches, such as lithography and micro-machining, have been extensively explored to mimic microvascular networks. Parthasarathy et al. [17] employed photolithography to generate cortical capillary patterns to cast planar Polydimethylsiloxane (PDMS) microfluidic phantoms with channel diameters ranging from 10 to 150 µm. Dong et al. [18] further extended this approach to replicate a cerebral vascular tree from a mouse brain image, while Luu et al. [19] demonstrated laser micro-machining to engrave channels into multilayer adhesive films. Additive manufacturing offers further control of channel geometry, scale, and branching topology. However, despite these technological improvements, several limitations remain concerning predefined designs, channel morphology, scalability, fabrication complexity, and cost.

A key limitation of all the above approaches is the necessity of an a priori design of the native vasculature model which restricts the natural variability present in real vascular systems. In addition, constructing networks spanning micron to millimeter scale vasculature typically requires multiple fabrication strategies, increasing fabrication difficulty and cost. Moreover, many approaches fail to replicate the smooth, circular cross-sections of native blood vessels, instead producing rectangular or stair-stepped geometries that alter flow dynamics as reported in [20]. Although some fabrication techniques are capable of producing circular channels, accurately replicating tortuosity and complex branching patterns, features critical to microvascular studies [21], remains challenging. Additionally, many phantoms are inherently planar, whereas native vasculature is three-dimensional and conforms to anatomical curvature, such as in the cerebral cortex or retina. Although some attempts have incorporated curvature, such as rigid conical molds in [22], these approaches lack flexibility and generalizability. Resin-based 3D-printed phantoms address some challenges but remain technically demanding, costly, and limited in scalability [23, 24].

To address these limitations, we introduce a lithography-free approach using the Lifted Hele-Shaw Cell (LHSC) technique. LHSC can naturally generate multiscale, fractal-like networks with smooth Gaussian cross-sections and progressive tapering of vessel diameters, which are characteristic of natural vasculature, without the need for predefined masks or serial patterning steps [25, 26]. While LHSC has been extensively explored in fluid dynamics for generating branched, fractal interfacial patterns reminiscent of natural networks like the blood brain barier and the lung tree [27, 28], its application as a direct fabrication strategy for tissue-relevant perfusion phantoms has not been realised. Building on our prior work [29], where LHSC was used to generate a fractal-like mask for phantom fabrication, we extend this approach to develop four distinct phantom designs. We further evaluate their functional relevance using laser speckle contrast imaging (LSCI), a widely used non-invasive optical modality for assessing superficial blood flow based on near-infrared light scattering. LSCI is particularly well suited for evaluating perfusion dynamics in vascular networks and has been extensively applied in neurology [30, 31], dermatology [32, 33], ophthalmology [34], and preclinical research [10, 35], where accurate characterization of microcirculatory flow is critical.

To establish the structural and functional similarity of the fabricated phantoms to native vasculature, we systematically refined the design and characterized the resulting networks by direct comparison with vascular morphology and perfusion measured in animal models using LSCI. In this work, we present the design and fabrication of a planar-tree phantom that captures the hierarchical branching and physiologically relevant flow distribution. The volumetric flow rate in the fabricated channels are compared to mouse cortical vasculature using generalized Murray’s law. To further improve anatomical relevance, we develop a guided morphology phantom incorporating a predefined anatomical guide and perform quantitative comparison with *in vivo* perfusion images. To move beyond planar structures, we introduce a flexible mold approach using LHSC that conforms to curved tissues such as cortex, retina, or skin. As a proof of concept, we fabricate a curved substrate phantom shaped after mouse somatosensory cortex. Finally, we establish the versatility of this technique across both animal and human vasculature by morphologically mimicking the radial organization of human retinal vasculature and use the Murray’s law to validate the flow–channel relationship by means of the angular LHSC method.

En route to validating the phantom using LSCI, we also note its potential application as a calibration platform for other speckle-based perfusion methods, including Laser Doppler Flowmetry (LDF) [36], Optical Coherence Tomography (OCT) [37], Diffuse Optical Spectroscopy/Tomography (DOS/DOT) [38], Diffuse Correlation Spectroscopy/Tomography (DCS/DCT) [39], Speckle Contrast Optical Spectroscopy/Tomography (SCOS/T) [40], Multi-speckle Diffuse Correlation Spectroscopy/Tomography (M-DCS/T) [41–43].

## 2 Results

### 2.1 Planar Tree Phantom

To establish a baseline physiologically inspired model, we first designed a vascular tree structure using the LHSC technique. The resulting structure comprises a central feeding vessel with symmetric, hierarchical bifurcations that taper and terminate at the periphery. The fabrication steps for the vascular tree mask using LHSC and the phantom are provided in the Methods section.

Fig. 1a(i) shows the vascular tree mask of dimensions 15 *mm×* 30 *mm*. It contains a central main channel with progressive bifurcations. White-light interferometry (MSA 500, Polytech, Germany) confirmed the hierarchical tapering of the channels, as shown in Fig. 1a(ii), where three representative branches corresponding to channels labeled as ‘A’, ‘B’ and ‘C’, exhibit heights of 122 *µm*, 59 *µm*, and 41 *µm* and widths of 830 *µm*, 240 *µm*, and 190 *µm*, respectively, each with a smooth Gaussian cross-section. The PDMS-based phantom obtained from this mask is shown in Fig. 1b(i). It includes a central inlet (green circle) and two peripheral outlets (yellow circles) to direct flow through the branching structure. The LSCI perfusion image in Fig. 1b(ii) demonstrates qualitatively that flow decreases with decreasing channel diameter, consistent with physiological vascular transport patterns [44].

**Fig. 1.**
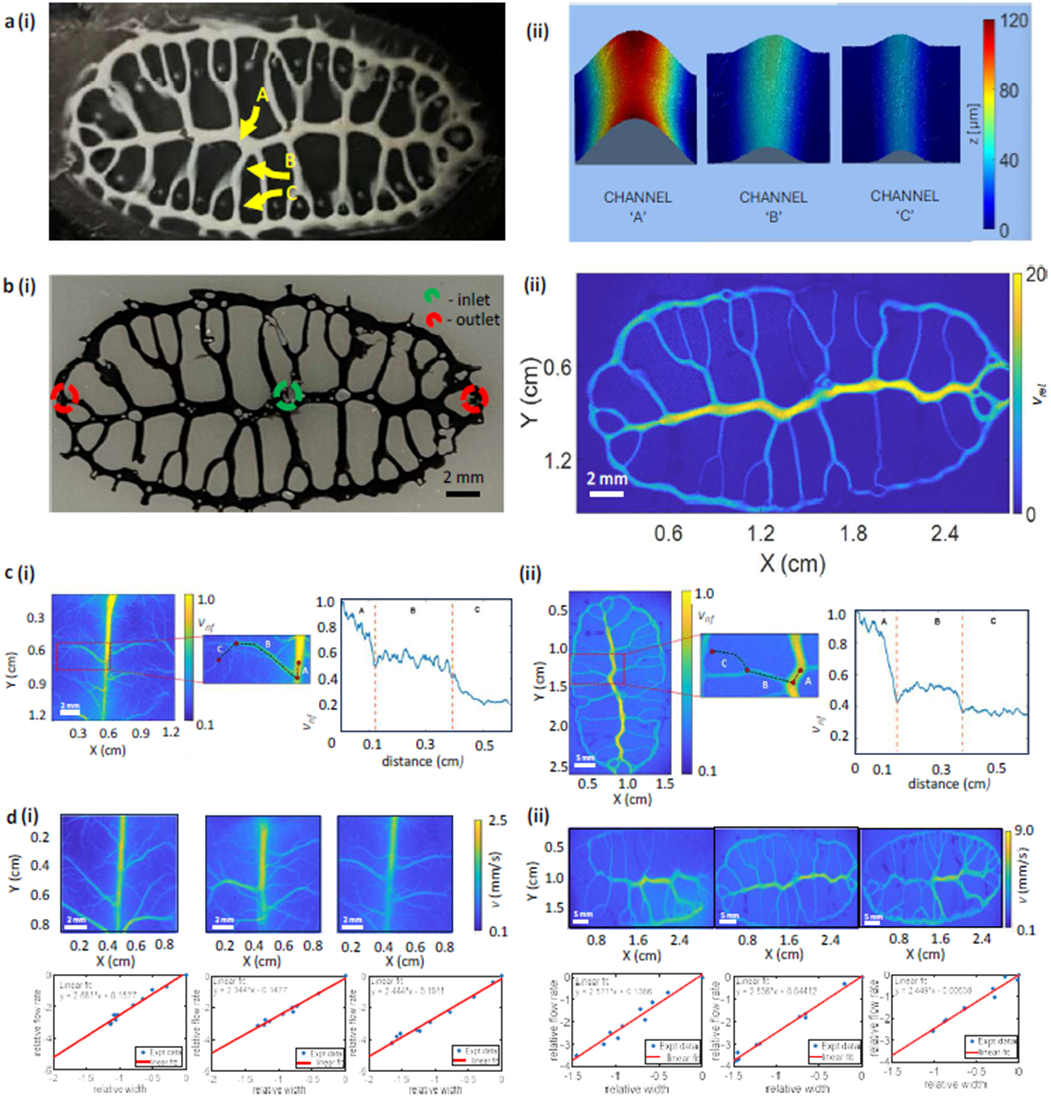
The planar-tree phantom. **a** (i) Planar-tree master mold fabricated using the Lifted Hele-Shaw Cell (LHSC) technique. (ii) Mask characterization by white-light interferometry showing the height, width, and surface profiles of channels A, B, and C (yellow arrows in **a(i)**). The channels exhibit a smooth Gaussian cross-sectional profile and a progressive decrease in height from A to C. **b** (i) Final PDMS-based phantom fabricated via soft lithography, showing the inlet (green circle) and outlets (red circles). TiO_2_ was mixed with PDMS as a scattering agent to mimic the optical scattering properties of biological tissue. The phantom channels were filled with ink for visualization. (ii) Relative flow velocity (*v*_*rel*_) map acquired using Laser Speckle Contrast Imaging (LSCI) (see Methods), showing perfusion within the phantom. Flow velocity is highest in larger-diameter vessels (e.g., the central vessel) and progressively decreases with decreasing vessel diameter. **c** Comparison of flow profiles between (i) the somatosensory cortex of a mouse brain and (ii) the planar-tree phantom. A representative vessel (red dotted line, inset) was selected from each perfusion image. The normalized flow (*v*_*nf*_) plotted along the vessel length demonstrates a similar flow distribution in both the mouse brain and phantom vessels. **d** Quantitative comparison between (i) three mouse brains and (ii) three planar-tree phantoms. Flow velocity (*v*) values from ten regions of interest (ROIs) and their corresponding vessel diameters are plotted on a log–log scale. Linear fitting was performed to estimate the Murray exponent, quantifying the similarity in the flow–diameter relationship between the phantom and *in vivo* mouse brain vasculature.

A quantitative comparison of perfusion in the phantom with *in vivo* behavior is provided in Fig. 1(c) and (d). Fig. 1c(i) and (ii) compares flow variation along vessels in the somatosensory cortex of the mouse brain and in the vascular phantom respectively. In both cases, flow decreases from regions A to C along branches of progressively smaller width. In Fig. 1d, flow values and their corresponding vessel diameters, obtained from ten distinct regions of interest (ROI), in each mouse and phantom perfusion maps are plotted. These measurements were subsequently analyzed to evaluate the scaling relationship between flow and vessel diameter according to Murray’s law. Across datasets from three mice, the Murray exponent was found to be 2.49 ±0.173, while three independently fabricated planar-tree phantoms yielded an exponent of 2.52 ±0.063, with no statistically significant difference between the two groups, which was determined using a two-sided t-test (p = 0.77). The relevant equations used to extract the flow information from LSCI data and to compute the Murray exponent are described in the Methods section.

### 2.2 Guided Morphology Phantom

Having validated the baseline vascular architecture, we next aimed to create phantoms that replicate specific anatomical vasculature rather than generic branching patterns. To do this, we developed a guided–fabrication approach, where the vasculature of interest served as an initial template for mask formation. Fabrication steps using LHSC are provided in the Methods section. Fig. 2a shows the workflow starting from a 0.8 *cm×* 0.8 *cm* region of the somatosensory cortex, including the Superior Sagittal Sinus (SSS) and associated branches. After skeletonization, the design was 3D-printed onto a quartz substrate. This 3D-printed design, known as anisotropy, directed the fluid flow in a specific direction during LHSC operation, enabling the repeated generation of guided masks. The final PDMS phantoms exhibit channels along the guide structure, along with additional spontaneous bifurcations resulting from inherent variability in the LHSC process.

**Fig. 2.**
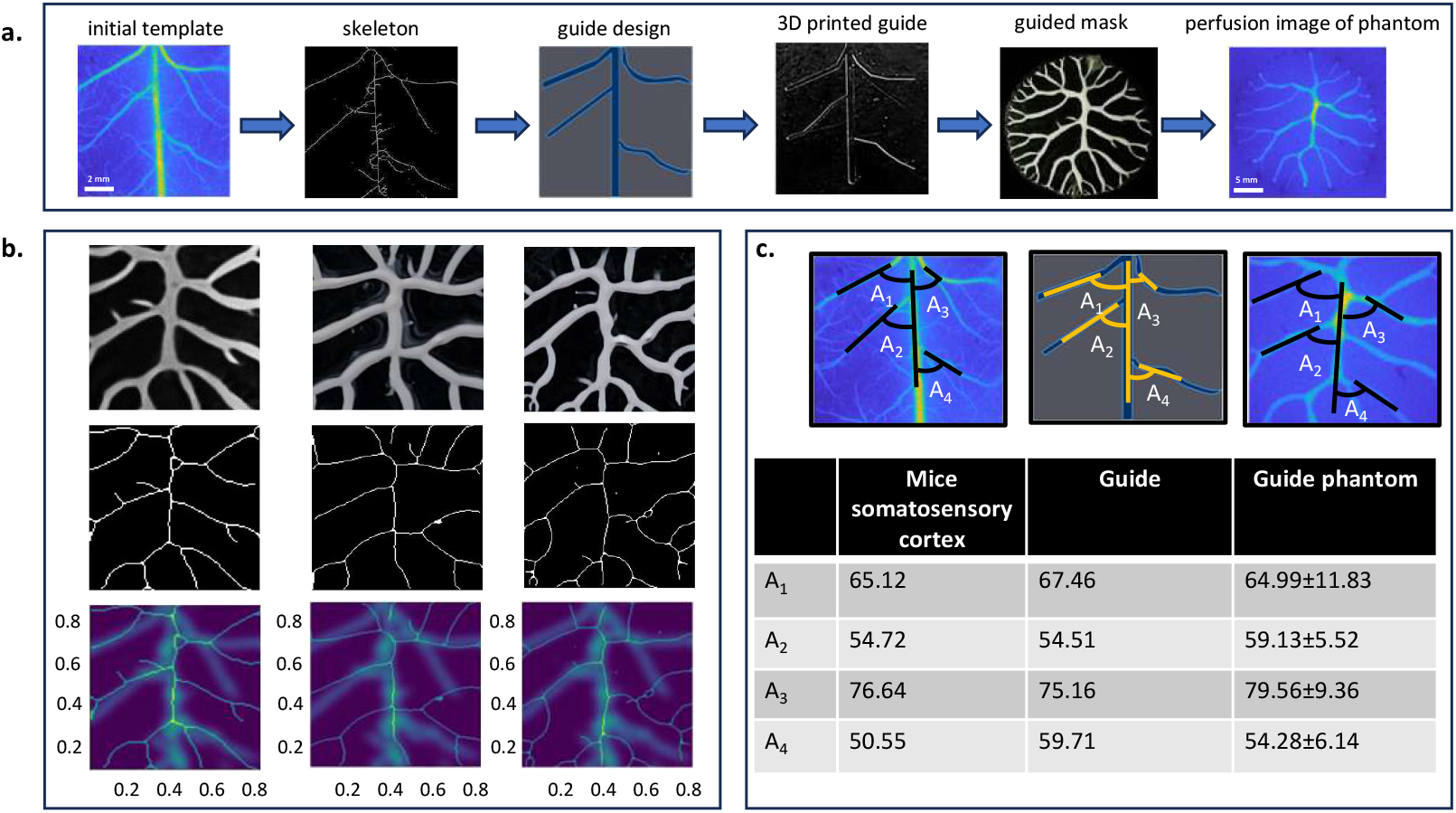
The guided morphology phantom. **a** Fabrication workflow of the guided-morphology phantom, starting from a perfusion map of a selected mouse-brain region used as the initial template, followed by guided mask design and soft-lithography fabrication of the final phantom. **b** Structural comparison between the initial template and three masks produced from the same 3D-printed guide, showing substantial spatial overlap of channel patterns with the original template. **c** Quantitative comparison of bifurcation angles across mouse-brain vasculature, the 3D-printed guide, and the guided-morphology phantom demonstrating close structural agreement between the fabricated phantoms and the original brain ROI.

Structural consistency across masks is shown in Fig. 2b, where three skeletonized masks overlay with a Gaussian-filtered skeleton of the original perfusion image (kernel: 90 *µm*, set by the minimum vessel diameter in the guide). Major features of the SSS align well with the *in vivo* structure, though bifurcation angle variability is observed.

The comparison of four SSS branches across multiple phantom replicas shown in Fig. 2c demonstrates that the guided morphology phantom reproduces both the anatomical geometry and natural morphological variability where the former is driven by intentional guidance and the latter emerges naturally due to the inherent randomness associated with the LHSC process.

### 2.3 Curved Substrate Phantom

To acheive more physiological relevance of the phantoms, we transitioned from planar to non-planar substrates for fabrication of the fractal-like masks. *in vivo* vascular networks, such as those in the cortex and retina, are distributed over intrinsically curved anatomical surfaces. Accordingly, we fabricated a curved substrate phantom using a non-planar mold to generate continuous vascular structures conforming to a three-dimensional geometry. The steps for the mask making and the subsequent phantom fabrication have been described under the Methods section.

Fig. 3a shows the phantom with ink-filled channels for visualization. For comparison, a 3D-printed mouse brain (Allen Brain Atlas, https://atlas.brain-map.org [45], dimension-matched to a 5-month-old mouse) is shown. Measurements from the chord and saggita (dotted lines) yielded a radius of curvature of 7.7 mm for the phantom and 6.2 mm (using Eq. 10) for the corresponding region of the mouse cortex (Fig. 3b). The LSCI perfusion map in Fig. 3c illustrates how curvature affects imaging, with regions away from the focal plane appearing increasingly defocused.

**Fig. 3.**
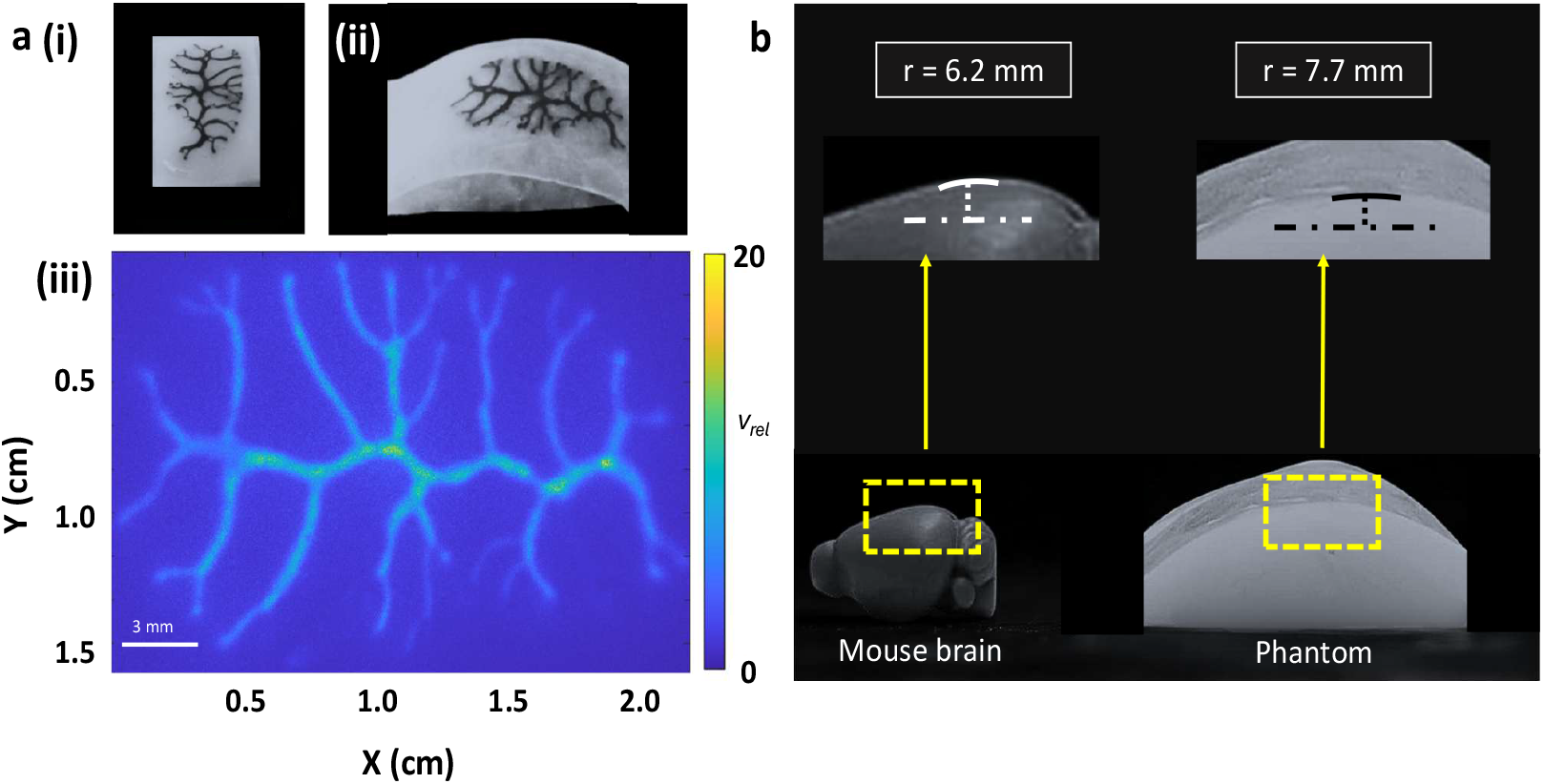
The curved substrate phantom. **a** (i) and (ii) visualisation of the channels in the curved substrate phantom filled with ink. (iii) perfusion map of the phantom obtained via LSCI. **b** saggital view of the 3D printed mouse brain (left) with dimensions to match that of a 5 month mouse and the curved substrate phantom (right) fabricated to mimic the curvature of the somatosensory cortex of the mouse brain (yellow box). Dotted lines indicate the chord (dot-dash) and the saggita (dot) drawn to estimate the radius of curvature (r).

### 2.4 Asymmetric Radial Network Phantom

Finally, to demonstrate adaptability across species and organs, we extended the approach to human vasculature by replicating the morphology of the retinal vascular network. Fig. 4a shows a white-light fundus image and its segmented representation [46]. Using LHSC with angular plate separation, two radial asymmetric masks were fabricated (Fig. 4b) from which the phantoms was made and the perfusion was visualized using LSCI (Fig. 4c). Five ROIs (width range: 150–350 *µm*) were analyzed, yielding Murray’s exponents of 2.75 and 2.73, closely matching reported human retinal values (2.74 for large arteries [47]), as shown in Fig. 4d.

**Fig. 4.**
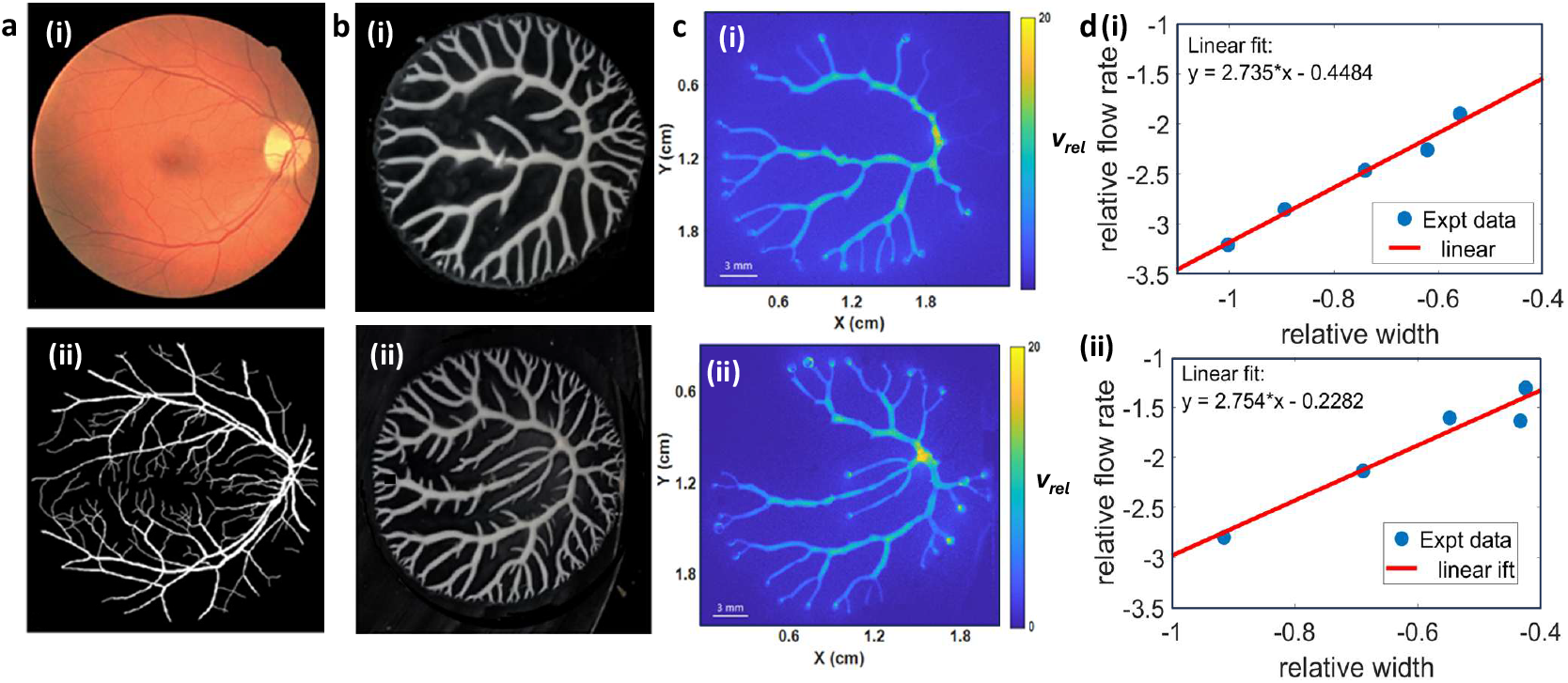
The asymmetric radial network phantom. **a** white light fundus image (top) and its segmented image (bottom) taken from [46]. **b** Masks made using LHSC with angular separation of plates to mimic the asymmetric radial structure of the retinal vessels as seen in (**a**). **c** perfusion images of the phantoms made using the masks in (**b**).[Phantom dimension = 15 mm x 15 mm; overall vessel diameter = 150-600 *µm*] **d** generalized Murray’s law analysis of the two phantoms for vessel diameters upto 350 *µm*.

## 3 Discussion

A central motivation of this study was to fabricate vascular phantoms that reflect the hierarchical, multiscale organization inherent to biological vasculature. Rather than imposing geometries a priori, we sought a fabrication approach in which such complexity could emerge naturally from the patterning process itself. The LHSC technique is well established for generating branching structures analogous to those found in natural systems, including snowflakes, leaf veins, and vascular networks. Its ability to spontaneously produce fractal-like networks spanning multiple length scales in a single step, combined with its experimental simplicity and low cost, made it particularly well suited for vascular phantom fabrication. These characteristics motivated our selection of LHSC as the underlying framework for our work.

Building on this framework, we introduced and validated a modular approach for fabricating anatomically inspired vascular phantoms. Beginning with a planar-tree design, we demonstrated that the phantom supports physiologically relevant flow behaviour, with measured flow–diameter relationships closely matching the predictions of generalized Murray’s law for mouse cortical vasculature. To further enhance the anatomical correctness, we incorporated an anatomical guidance in the mask making method and quantitatively compared the resulting phantom to *in vivo* perfusion maps. Extending beyond planar fabrication, we developed a flexible mold approach to generate phantoms that conform to physiologically relevant three-dimensional curvature (cortical, ocular or dermal), which in this case we demonstrated by using the geometry of mouse somatosensory cortex. Finally, the method was generalized to human vascular morphology by recreating a radial vascular organization of the retina by angular LHSC method and validated the flow–diameter relationship using the generalized Murray’s law.

A key strength of the proposed phantoms is the fractal-like self-organization inherent to the LHSC process, which enables spontaneous and scalable formation of multiscale vascular networks with channel widths spanning from a few microns to several hundred microns. In fact, several masks developed in this work, and their corresponding PDMS molds, contained channels with widths below 100 *µm*. However, these finer channels did not exhibit observable perfusion.

This limitation arises from two primary factors. First, the minimum diameter of commonly available PDMS punch tools is approximately 300 *µm*, which prevented the creation of outlets for smaller peripheral channels. As a result, although these fine structures were successfully replicated in the PDMS, the absence of dedicated outlets restricted flow in these regions. More precise outlet fabrication could be achieved using microscope-assisted micro-drilling approaches, such as CNC-based drilling or pulsed laser ablation techniques [48]. Second, the spatial resolution of the widefield LSCI system used in this study was approximately 100 *µm*, which limits the reliable visualization of perfusion in channels below this scale. Imaging flow in smaller vessels would require higher-resolution microscopic perfusion imaging systems [49].

We emphasize that a core objective of this work was to demonstrate that anatomically realistic perfusion phantoms can be fabricated using commonly available tools and materials. Within these practical constraints, this work demonstrates that anatomically realistic vascular organization and physiologically relevant flow behaviour can be achieved using commonly available tools and materials, while still retaining flexibility for further scaling and refinement. By modulating parameters like the fluid film thickness after squeezing and the velocity of separation of the plates [25] masks with a broader range of channel widths can be generated. Such multi-scale phantoms could serve as calibration standards for both microscopic and macroscopic imaging modalities.

Due to the above mentioned bottlenecks, we have noticed that compared to mouse cortical vasculature, our planar tree phantoms exhibit larger vessel diameters and higher flow rates as seen in Fig. 1. However, the Murray exponents obtained from the phantoms align well with the reported biological values with a variance smaller than that obtained in the mouse brain case (for animal = 2.49 ±0.173; for phantom = 2.52 ±0.063; p = 0.77) [50–53]. This shows the physiological relevance, reproducibility and robustness of the phantom. In fact, with such realistic perfusion models the dependency on animal-based experimentation could be minimized leading to more sustainable and ethical preclinical research practices.

The guided morphology phantom demonstrates a rapid and effective approach for structurally replicating vasculature from a specific anatomical region, with acceptable variance as shown in Fig. 2c. This tolerance can be further improved with more accurate guide design and tighter control of mask-generation parameters, as discussed earlier. Building on this concept, the curved substrate phantom increases anatomical relevance by incorporating surface geometry. While the current implementation mimics the uniform curvature of the mouse somatosensory cortex, more complex and non-uniform topographies representative of real tissue surfaces can be generated. The flexible mask used for curved fabrication can be made from any stretchable substrate capable of conforming to three-dimensional surfaces, and because the mask remains fluid prior to curing, its geometry remains preserved during deformation. An illustration of the same is shown in Fig. 5 where the flexible mask with a planar vascular-tree design has been molded to a 3D printed brain showing the potential of this technique to eventually generate realistic perfusion phantoms with complex curvatures. Therefore, both the guided and curved approaches enhance anatomical realism and enable application-specific customization.

**Fig. 5.**
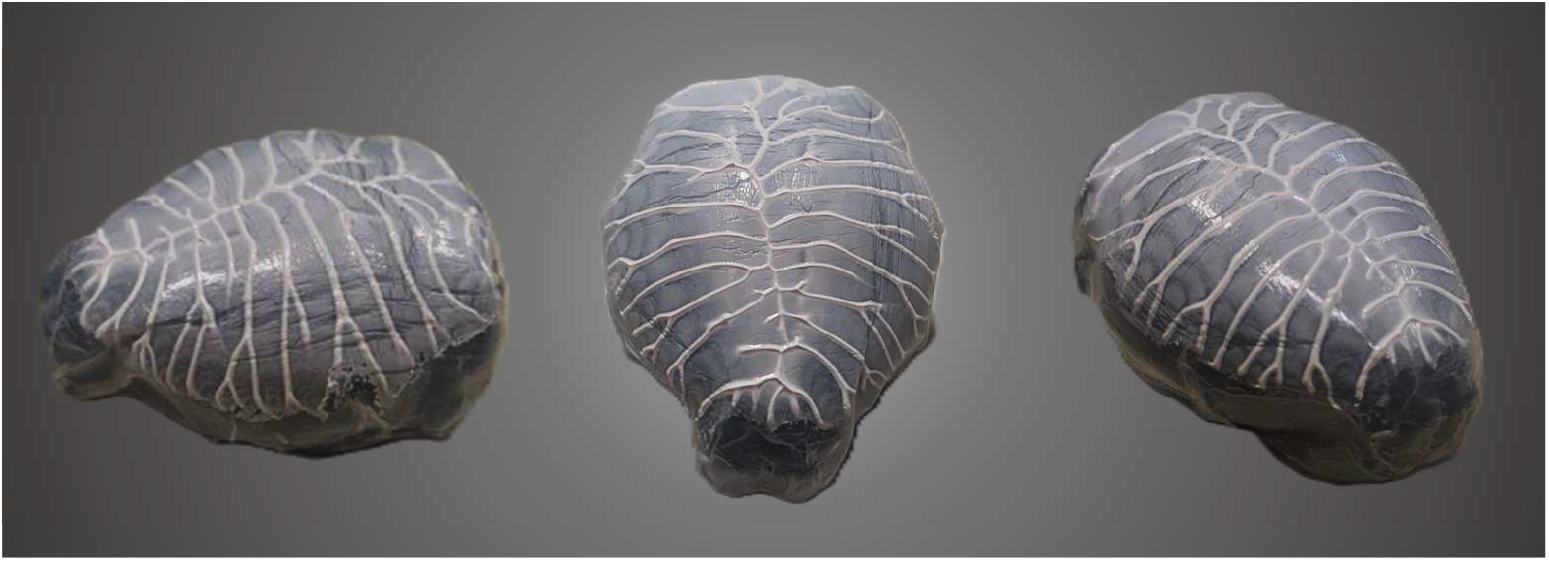
Illustration of a 3D brain phantom. Implementation of the curved-mask fabrication technique to obtain the overall anisotropic curvature of brain. Here a flexible vascular tree mask has been conformed against a 3D printed mouse brain to reproduce the cortical geometry.

Curvature incorporation is particularly relevant for optical modalities where surface topography introduces distortions or depth-related scaling errors [54, 55]. This is especially pertinent for tomographic techniques, in which reconstruction algorithms often rely on simplified acquisition geometries that implicitly assume planar surfaces [10, 40, 43]. In practice, imaging is frequently performed over anatomically curved boundaries, leading to deviations in illumination and detection geometry, variations in the effective source–detector separation, and changes in the local laser spot profile during scanning. These effects propagate into the forward model and can introduce systematic errors in the reconstructed images if unaccounted for. Curved vascular phantoms therefore provide a controlled platform for developing and validating correction strategies and reconstruction algorithms for non-planar imaging geometries.

The modularity of the four phantom configurations presented here, planar tree, guided morphology, curved substrate, and asymmetric radial network, allows them to be combined as needed. For instance, an anatomically guided vascular mask can be fabricated on a flexible substrate and subsequently shaped to define regional curvature. Similarly, a retinal-morphology phantom could be formed on a hemispherical mold to better emulate the globe geometry for OCT or adaptive optics device testing. Furthermore, the PDMS substrate may be substituted with alternative TMMs like agarose or polyvinyl alcohol (PVA) cryogel, to extend compatibility to MRI, ultrasound, or other perfusion imaging systems.

Beyond perfusion imaging, such biomimetic models of vascular systems hold value in endovascular catheterization techniques and instruments [56], microsurgical simulation [57], vascular pathology modeling, and drug transport studies including tumor microenvironment research [58, 59]. As the vascular structures are not pre-designed but emerge from the experimental conditions such as fluid properties and flow parameters, this phantom offers a versatile testbed for biological, clinical, and engineering research domains where controlled, repeatable, yet physiologically plausible vasculature is required.

## 4 Methods

### Animal preparation

The experimental procedures used in this work was approved by the Institutional Animal Ethics Committee (IAEC) at IISER Pune, and the Committee for the Control and Supervision of Experiments on Animals (CCSEA), Government of India (animal facility CCSEA registration number 1496/GO/ReBi/S/11/CCSEA). The usage of animals in this manuscript was approved under protocol number IISER-Pune/IAEC/2019-2/04. Three C57BL/6 mice, aged 4-5 months and weighing 25-30g, with skull removed over the somatosensory cortex of the brain were used for the study. Common surgical procedures were followed in preparaing the animals for *in vivo* imaging, as described before [60, 61]. The mice were initially anesthetized with ketamine and xylazine and then imaged using our in-house developed small animal imaging platform [10].

### Lifted Hele-Shaw Cell technique (LHSC)

When a non-Newtonian fluid confined between two plates is separated, branching or fractal-like features form at the fluid–air interface due to Saffman–Taylor instability, which occurs when a fluid of low viscosity dynamically displaces a fluid of high viscosity. This process forms the basis of the lifted Hele-Shaw Cell (LHSC) technique [62, 63]. Islam et al.[25] demonstrated that such structures can be retained permanently by using a ceramic precursor fluid that cures upon UV exposure. During lifting, air acts as the low-viscosity displacing phase, while the ceramic suspension fluid behaves as the more viscous phase, generating mirror-symmetric patterns on both plates. After curing, one of the plates serves as the solid master mold (Fig. 6a). The non-Newtonian ceramic suspension photo polymer solution consists of monomer HDDA (1,6-Heaxanediol diacrylate), Alumina powder (mean particle size 0.5 micron), photo-initiator Benzoin Ethyl Ether (BEE) 4 wt. % with respective to weight of monomer and surfactant Phosphate Ester 2.5 wt. % with respective weight of alumina powder.

**Fig. 6.**
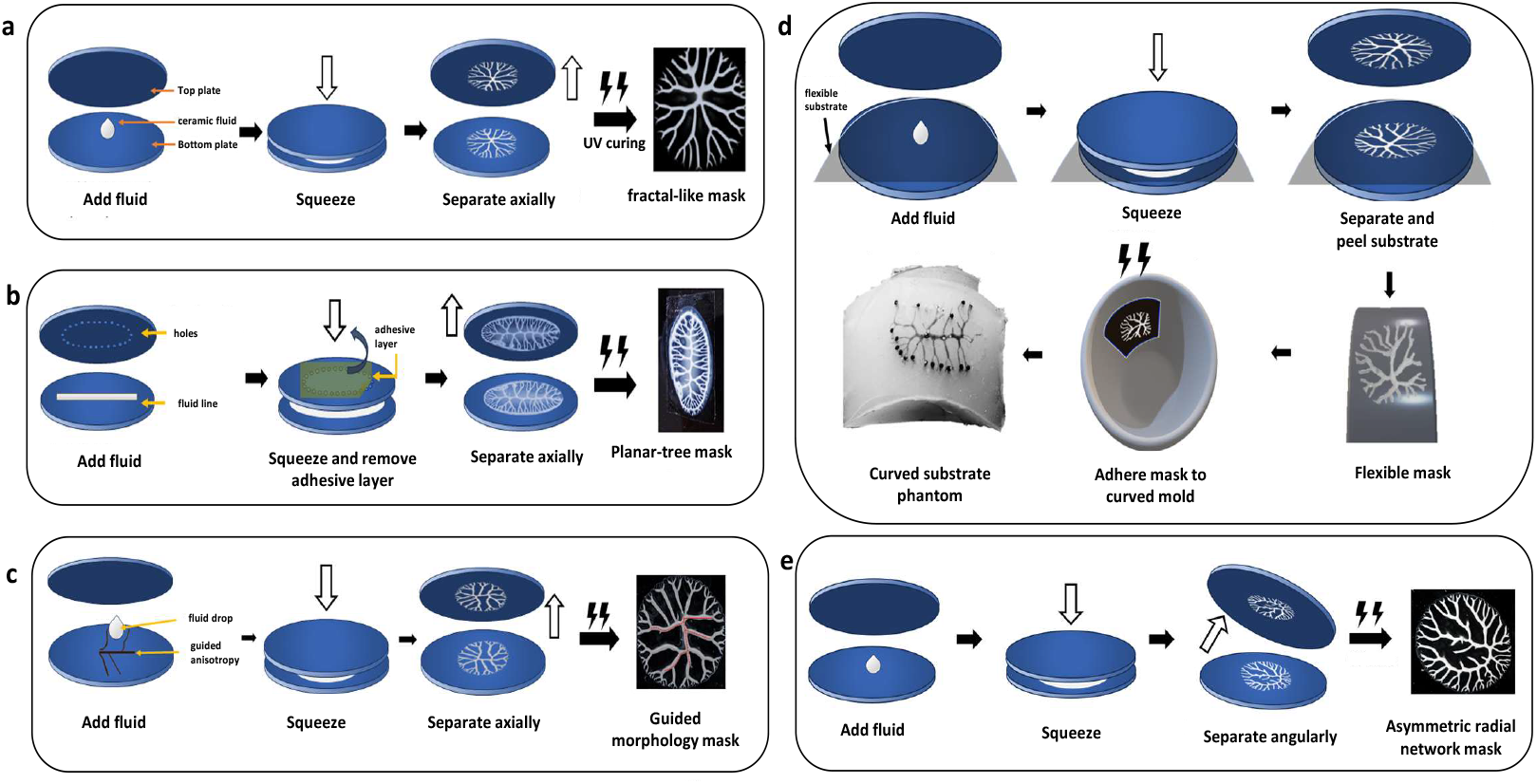
Schematic outlining the steps of mask creation. **a** fractal-like mask obtained using conventional LHSC **b** generic planar tree mask **c** guided morphology mask **d** curved substrate phantom creation with a flexible and stretchable mask **e** asymmetric radial network mask

### Planar tree phantom

To generate a physiologically inspired branching pattern, the LHSC method was adapted with multiport arrangement that controls the propagation of the Saffman–Taylor fingers such that all branches remain connected at the boundary [26] (Fig. 6b). Using this a vascular tree structure was created which is defined as a single primary channel that progressively bifurcates into smaller branches.

A linear volume of ceramic precursor was first deposited on a substrate to form the main trunk before squeezing the plates. The top plate included 30 equidistant holes (diameter 0.03 cm), arranged elliptically (semi-major axis: 1.25 cm, semi-minor axis: 0.75 cm). Upon separation, the precursor expanded preferentially toward these ports, producing a controlled branching morphology. The extent of branching and channel width were tuned by varying precursor volume and plate separation velocity.

After curing, the mold was used for soft lithography in PDMS. Titanium dioxide (TiO_2_) was incorporated at 1.8 mg/g PDMS [17] to achieve a scattering coefficient of ~ 10 cm^−1^, representative of biological tissue. Following curing, inlet and outlet ports were punched. Due to the connected outer branches, only one inlet and two outlets were required.

### Guided Morphology Phantom

To reproduce anatomically accurate vasculature rather than generic branching, a guided anisotropy method was used. Major vessels from a selected region of interest were printed onto a substrate via 3D microfabrication, and this patterned anisotropy served as a flow guide during LHSC [64].

For this work, a segment of the Superior Sagittal Sinus (SSS) from the lambda–bregma region of a three-month-old C57BL/6 mouse was used. The input perfusion image was binarized and skeletonized using the `skeletonize` function from the scikit-image Python library [65]. The original vessel diameters were preserved and assigned an empirically chosen uniform height of 50 *µm*. The corresponding mask is shown in Fig. 6c.

During the LHSC process, the precursor fluid preferentially adhered to the printed anisotropic lines, allowing the final cured mask to contain both the anatomically guided vessels and additional stochastic fractal branches. This method therefore enables fabrication of phantoms containing user-defined vasculature with controlled variability in the branching patterns.

### Curved Substrate Phantom

To demonstrate applicability to non-planar biological geometries, the fabrication approach was extended to curved substrates.

For the mask creation a flexible substrate (PVC or PDMS sheet) was temporarily bonded to the lower plate during LHSC (Fig. 6d). After creating the structure, the uncured mask was peeled from the substrate and conformed to the desired curved surface. Because the ceramic precursor remained fluid prior to curing, it redistributed to maintain smooth channel morphology despite mechanical stretching. UV exposure was then used to cure the mask and form a curved master mold.

A PDMS–TiO_2_ mixture was then poured into the curved mold, cured, and subsequently bonded to a thin PDMS layer after punching inlet and outlet ports. The resulting phantom surface’s curvature corresponds to the negative of the mold curvature. Therefore, producing a phantom with a desired anatomical radius requires designing the mold with the inverse geometry. The full fabrication sequence and final curved substrate phantom are shown in Fig. 6d

### Asymmetric Radial Network Phantom

To demonstrate scalability to human vasculature, the technique was applied to create a phantom representing the coronal branching of retinal vessels. Unlike previous setups that used axial lifting, this version employed angular plate separation to mimic radial branching originating from the optic disc. The final structure (Fig. 6e) features vessel branches radiating outward in an arc, consistent with retinal anatomy.

### Validation using Laser Speckle Contrast Imaging (LSCI)

The flow validation in these phantoms was carried out using LSCI as shown in Fig. 7. A laser source of 785 nm (Thorlabs L780P090) with beam shaping optics was used for uniform illumination of the sample surface. The backscattered rays from the sample were focused and imaged using an sCMOS camera (Photometric Prime BSI) with a 50 mm objective lens (Edmund Optics) [43] resulting in a field of view of 30 mm x 30 mm covering full sensor area (2048 x 2048 pixels). The inlet of the phantom was connected to a syringe pump for pumping intralipid through its channels.

**Fig. 7.**
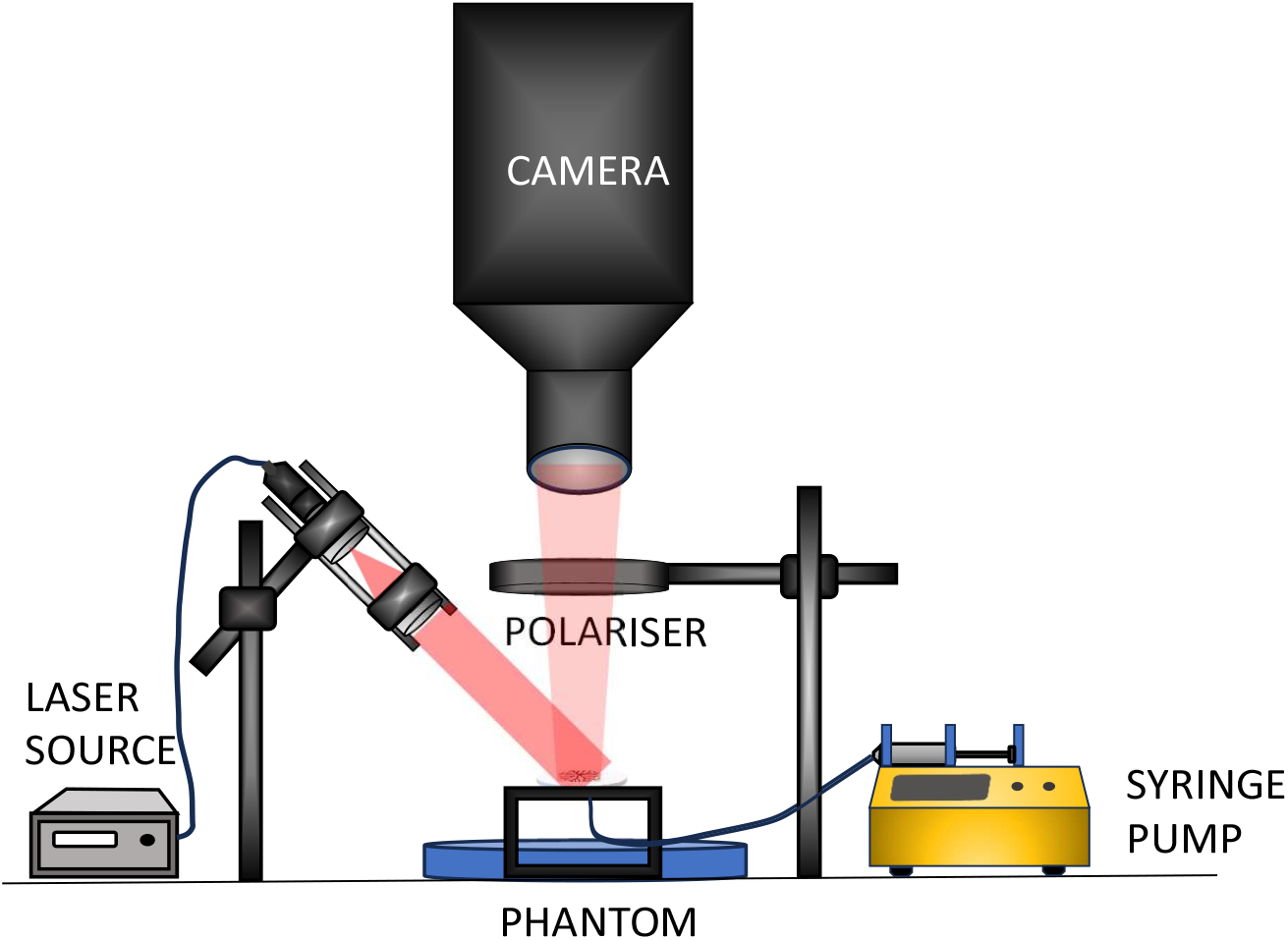
Schematic of LSCI experimental setup for validation of the fabricated phantom. A 785-nm laser beam is expanded and directed onto the phantom, and the backscattered light is imaged onto a camera. A polariser in the detection path suppresses specular reflections from the phantom surface. Intralipid is infused into the phantom at controlled flow rates using a syringe pump, during which raw speckle images are recorded and processed to generate speckle-contrast maps.

Multi-exposure images were acquired where fifteen logarithmically spaced exposures (*T*) in the range 50-3000 *µs* with 100 images per exposure were captured. The raw intensity data was processed to obtain the speckle contrast (*κ*(*T*)) for a given exposure time (*T*) at every pixel temporally, which is given as –

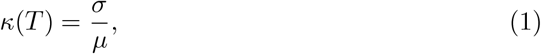

where *σ* and *µ* are temporal pixel-wise standard deviation and mean intensity, respectively. The multi-exposure speckle contrast was fitted to Eq. 2 to extract the decorrelation time *τ*_*c*_, inversely proportional to flow velocity *v* [10, 66, 67]:

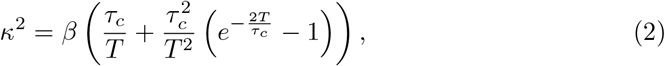

where *β* is a constant that depends on the collection optics of the experimental setup. Its value was determined to be 0.6 using a solid PDMS-based phantom and assumed constant throughout the experiment. The flow velocity was then determined as –

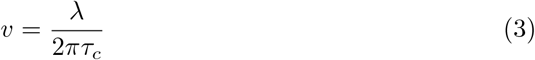

where *λ* is the wavelength of the incident light (*λ* = 785 nm).

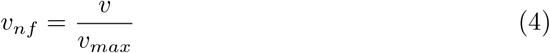

Normalized velocity and relative velocity maps were computed using Eqs. 4 and 5.

where *v*_*max*_ is the maximum velocity in the channel.

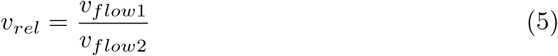

where *v*_*flow*1_ and *v*_*flow*2_ are velocities obtained corresponding to two different flow rates in the channel.

### Murray’s law

The quantification of flow in the fabricated channels was compared with *in vivo* mouse perfusion data using Murray’s law. According to Murray’s law, vascular branching follows an energy-minimizing principle that balances the metabolic cost of maintaining vessels with the pumping power required to drive blood flow. Under this condition, the volumetric flow rate (*Q*) scales with vessel diameter (*d*) as

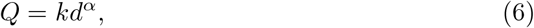

where *α* is the Murray exponent and *k* is a proportionality constant. For Newtonian fluids, *α* = 3, whereas for non-Newtonian fluids such as blood, *α* deviates from this ideal value [68, 69].

Dividing both sides of Eq. 6 by the maximum flow rate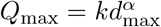 yields

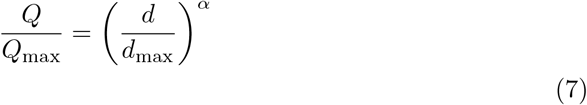

Defining the normalized flow rate *Q*_norm_ = *Q/Q*_max_ and normalized diameter *d*_norm_ = *d/d*_max_ and taking the logarithm of both sides gives,

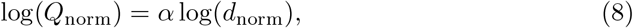

such that a log–log plot of *Q*_norm_ versus *d*_norm_ yields a straight line with slope *α*. For each phantom or animal perfusion map obtained using LSCI, mulitple ROIs were selected from channels or vessels spanning a range of diameters. The volumetric flow rate, *Q*, in each ROI was calculated as

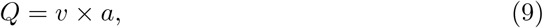

where *v* is the flow velocity obtained from Eq. 3 and *a* is the cross-sectional area. A circular cross-section was assumed for *in vivo* vessels (*a* = *πd*^2^*/*4) and a half-circular cross-section was assumed for the fabricated channels (*a* = *πd*^2^*/*8).

The resulting flow rates and corresponding diameters were normalized and plotted on a log–log scale. As indicated by Eq. 8, the Murray exponent *α* was obtained as the slope of a linear fit to this relationship.

### Radius of Curvature Measurement

To quantify the curvature of the curved substrate phantom, the radius of curvature (*R*) was estimated by assuming a circular arc over the curved surface as shown in Fig. 8. A chord of length *L* and a line perpendicular from the chord to the arc called the sagitta of length *h* are drawn. By using standard geometrical relations for circles, we get -After curing, the mold was used for soft lithography in PDMS

**Fig. 8.**
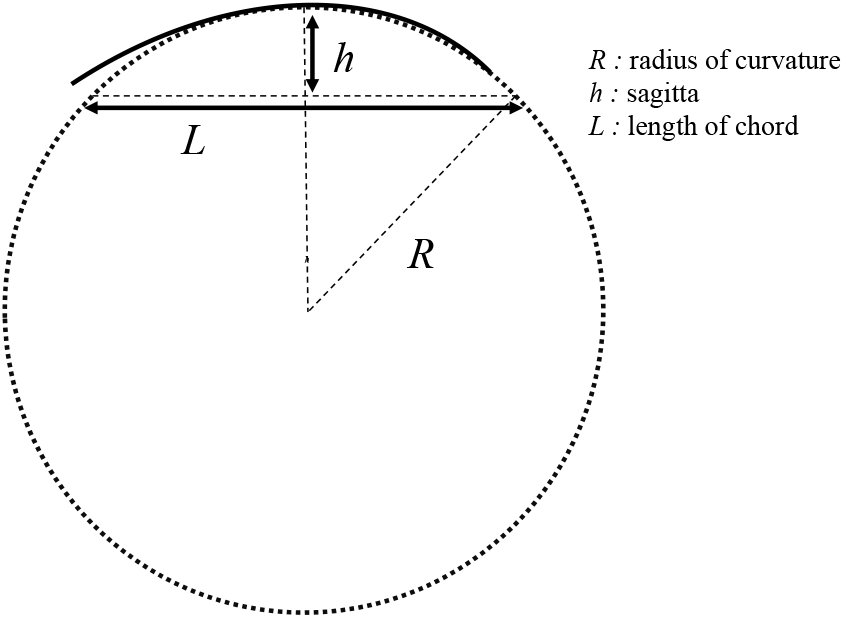
Determination of the radius of curvature. Schematic illustrating the estimation of the radius of curvature (R) of a curved surface from the measured chord length (L) and sagitta (h)

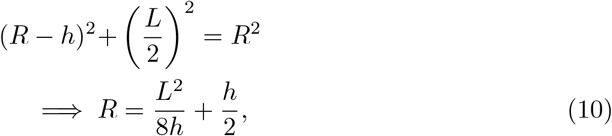

## Code, Data, and Materials Availability

The code and data utilized in this study are available from the authors upon reasonable request.

## Acknowledgments

This research was funded by Wadhwani Research Centre of Bioengineering, Indian Institute of Technology Bombay, and Biotechnology Industry Research Assistance Council (BIRAC, Government of India under the AIR-PACE scheme).

## Author contributions

S.D., M.R., S.S., R.P., P.S.G. and H.M.V. designed research; S.D. and M.R. performed research; S.D.M. and N.M.A. have performed the animal surgeries and contributed to *in vivo* data acquisition and consequent animal data validation; S.D. and H.M.V. analyzed data; and S.D, M.R., P.S.G. and H.V. wrote the paper.

